# Towards a More Objective and High-throughput Spheroid Invasion Assay Quantification Method

**DOI:** 10.1101/2024.06.27.600893

**Authors:** Rozanne W. Mungai, Roger J. Hartman, Grace E. Jolin, Kevin W. Piskorowski, Kristen L. Billiar

**Affiliations:** Department of Biomedical Engineering, Worcester Polytechnic Institute, Worcester, MA, USA 01605; independent researcher

**Author notes:** Correspondence to: Kristen L. Billiar Biomedical Engineering Department Worcester Polytechnic Institute 100 Institute Road Worcester, MA 01609.

## Abstract

Multicellular spheroids embedded in 3D hydrogels are prominent *in vitro* models for 3D cell invasion. Yet, quantification methods for spheroid cell invasion that are high□throughput, objective and accessible are still lacking. Variations in spheroid sizes and the shapes of the cells within render it difficult to objectively assess invasion extent. The goal of this work is to develop a high-throughput quantification method of cell invasion into 3D matrices that minimizes sensitivity to initial spheroid size and cell spreading and provides precise integrative directionally-dependent metrics of invasion. By analyzing images of fluorescent cell nuclei, invasion metrics are automatically calculated at the pixel level. The initial spheroid boundary is segmented and automated calculations of the nuclear pixel distances from the initial boundary are used to compute common invasion metrics (i.e., the change in invasion area, mean distance) for the same spheroid at a later timepoint. We also introduce the area moment of inertia as an integrative metric of cell invasion that considers the invasion area as well as the pixel distances from the initial spheroid boundary. Further, we show that principal component analysis can be used to quantify the directional influence of a stimuli to invasion (e.g., due to a chemotactic gradient or contact guidance). To demonstrate the power of the analysis for cell types with different invasive potentials and the utility of this method for a variety of biological applications, the method is used to analyze the invasiveness of five different cell types. In all, implementation of this high□throughput quantification method results in consistent and objective analysis of 3D multicellular spheroid invasion. We provide the analysis code in both MATLAB and Python languages as well as a GUI for ease of use for researchers with a range of computer programming skills and for applications in a variety of biological research areas such as wound healing and cancer metastasis.

## INTRODUCTION

Cell invasion and migration are driving factors for various biological events such as development [1,2], wound healing [3–6], tumor metastasis [7–9] and host cell infiltration of implanted scaffolds [10–13]. The effects of soluble and immobilized chemical gradients on cell migration on 2D surfaces and infiltration into 3D matrices and tissues is well established [9,14]; however, it is also becoming clear that mechanical cues play a large role in cell invasion and migration [7,15]. Cells move in response to physical cues such as stiffness gradients [9,15,16], contact guidance [9,15,17,18] and mechanical restraint [15,19–29].

*In vitro* models consisting of multicellular spheroids embedded in ECM are commonly used for investigating the mechanisms of cell invasion into extracellular matrices (ECM) in a variety of diseases [30–36]. We use the term cell invasion to refer to the 3-dimensional spread of cells in the ECM [30,37] which can occur by both cell proliferation and cell migration. Low contrast phase imaging is often used to capture images of cell invasion as it accommodates live-imaging. However, phase images of spheroid invasion are difficult to analyze with common automated software. Thus, this method often requires manual tracing of cell boundaries in programs such as ImageJ/FIJI to quantify the extent of invasion via measuring the distances and areas of migrating cells [8,30,38–44]. This manual tracing is time consuming and introduces subjectivity and possible technical error. Alternatively, high-contrast fluorescent images are amenable to automated quantification using common analysis software which allows for more objective high-throughput analysis. However, fluorescent images are less accommodating for live imaging as they require transfection of fluorescent proteins [37] or adding live cell stains which do not give a strong signal-to-noise ratio [45]. To overcome this limitation, clear high contrast fluorescence images can be captured by fixing and staining samples at the experiment endpoint using common accessible dyes. After imaging, the cell invasion can be quantified (such as with migration tracks [8,46]) using image analysis software or custom programs, and invasion metrics can be calculated.

Quantification metrics for cell invasion into 3D matrices commonly involve the overall invasion area and/or the distance travelled by cells at a given timepoint [30,34,37–39,42,44,47–49]. The invasion area is used to represent the number of invaded cells [30,34,37–39,42,44,49–51], and the distance travelled is often reported as either the mean or maximum distance travelled by cells [8,30,42,49,52]. Yet, these metrics are sensitive to changes in cell size and shape that occur as cells migrate through the matrix [53,54]. In the case of multicellular spheroid *in vitro* models, when invasion area and distance travelled are calculated without taking into account the initial spheroid boundary, these metrics are also sensitive to differences in size and shape of the spheroids. A metric that takes into account both the invasion area and distance travelled is especially useful, such as the migration index published by Liu *et al.* [30], as it informs both how many and how far cells are invading. The types of quantification metrics that can be calculated are limited by the methods used to capture and quantify images, especially if a common or microscope-based software is used. Therefore, there is a need for an easy-to-implement method and corresponding computational analysis tool that is precise and objective, offers flexibility in quantification, and is simple to use for a researcher unfamiliar with programming.

Here we present a high-throughput automated quantification method of cell invasion into 3D matrices using a complementary pixel-based method to segmentation methods. Multicellular spheroids embedded in collagen hydrogels are live-stained for cell nuclei, which are easy to image and are more resistant to shape changes during invasion than the cell body, and the distribution of nuclear pixels is quantified. Live fluorescent imaging allows for precise segmentation of the initial spheroid boundary, thus minimizing the effects of spheroid initial size and shape. Given binarized images, automated calculations of the invasion area as well as distances and angles of nuclear pixels from the boundary are calculated; these metrics are relatively insensitive to initial spheroid size. We introduce the area moment of inertia, which takes into account both the area and distances of nuclear pixels of invading cells, as an integrative metric for the overall extent of invasion. In combination with principal component analysis, the area moment of inertia metric is also used to quantify invasion directionality in response to anisotropic mechanical constraint. Five different cell types are included to demonstrate the applicability of the analysis for cell types with different invasive potentials, and we provide an open-source graphical interface and the detailed code in both MATLAB and Python languages for ease of adoption by researchers with a range of computer programming skills.

## MATERIALS AND METHODS

### Ethics

No experiments were performed on humans or animals for this study; therefore, ethics approval is not required. Freshly isolated cells used in this study were harvested from animal (pig) tissue while all other cells, including human cells, were obtained as a gift.

### Cell types and culture conditions

Cell spheroids of various cell types were embedded in collagen hydrogels and allowed to invade the matrix for two days. The cell types were chosen to span mesenchymal and epithelial cell type classifications as they are known to be motile and immotile respectively. Valvular interstitial cells (VIC), dermal fibroblasts and smooth muscle cells (SMC) are the mesenchymal cell types studied to highlight differences in invasive potential within the group and were compared to epithelial RPE-1 cells. PC9 cells were also included as we are interested in how a cancerous epithelial cell type (having undergone the endothelial-to-mesenchymal transformation) compares to non-cancerous mesenchymal cell types.

Porcine aortic VICs were isolated from fresh pig hearts obtained from a local abattoir (Blood Farm, Groton, MA), and the VICs were isolated within three hours as per published protocols [20] and used in experiments between passages 2-8. Neonatal human dermal fibroblasts (NHF), originally harvested from de-identified donated male foreskin tissue, were obtained as a gift from Dr. George Pins (Worcester Polytechnic Institute) and used in experiments between passages 6-12. Immortalized WKY 3M-22 male rat aortic SMCs were obtained as a gift from Dr. Marsha Rolle (Worcester Polytechnic Institute). The immortalized female human retinal pigment epithelial cell line (RPE-1) was obtained as a gift from Dr. Amity Manning (Worcester Polytechnic Institute). The same culture medium base formulation and conditions were used for all of the above cell types: DMEM (Gibco) supplemented with 10% v/v fetal bovine serum and 1% v/v antibiotic-antimycotic (Gibco). For the SMCs, 1% v/v MEM non-essential amino acids (Gibco) were added to maintain a growth phenotype. The PC9 human lung adenocarcinoma cell line was also obtained as a gift from Dr. Amity Manning and were cultured in RPMI-1640 (GenClone, Genessee Scientific) supplemented with 10% v/v fetal bovine serum and 1% v/v antibiotic-antimycotic (Gibco). The identity of the PC9 cancer cells have been previously confirmed by STR analysis [55]. The cell cultures were maintained at 37°C in a humidified 5% CO_2_-containing incubator.

### Generation of multicellular spheroids

3D spheroids were generated for each cell type as in our previous work [56]. Briefly, the cells were trypsinized, resuspended, and stained with 5μg/mL Hoechst 33342 (Invitrogen) for 10 minutes at 37°C for visualization of cell nuclei. The stain was then diluted, and the cells were resuspended at a cell concentration of one million cells per 200 μL. Non-adherent agarose microwells were generated by pipetting a pre-warmed 2% w/v agarose solution (MilliporeSigma), made in DMEM, into negative molds with 500 μm diameter wells (24-96 or 12-256; Microtissues®), and the agarose was allowed to solidify. The stained cell suspension was seeded into the prepared agarose microwells, and the cells were allowed to settle into spheroids for one day (Supplementary Fig. 1) before embedding them into collagen hydrogels.

### Embedding of multicellular spheroids into collagen gels

The multicellular spheroids were harvested from the microwells and suspended in media. Cooled rat tail collagen type I (Advanced Biomatrix, RatCol®) was then mixed with neutralization solution following manufacturer’s recommendations to make collagen hydrogels. The spheroid suspension was mixed into the collagen solution for ∼ 20 spheroids/mL in a final collagen concentration of 2 mg/mL. Next, the solution was poured into well plates and allowed to gel for 30 minutes at 37°C. The well plates used were either a tissue culture plastic 6-well plate for maximizing gel volume, or a flexible silicone well plate with small side posts (CellScale MechanoCulture FX™) to restrict gel compaction (4×4 array of 8 mm x 8 mm square wells).

Fresh medium was added over the samples, and the samples were imaged (Day 0) as described below. After imaging, the samples were returned to 37°C in a humidified 10% CO_2_-containing incubator and allowed to culture for two days (Supplementary Fig. 2). The timepoint of two days was chosen to allow sufficient time for cells to invade into the matrix while minimizing migration past the camera field of view.

### Staining and Imaging

The extent of cell invasion and migration was assessed by capturing images of the cell nuclei as 10µm- spaced z-stack slices through the thickness of the spheroids using a Keyence BZ-X810 fluorescence microscope (6.1µm DOF, High Resolution setting: 6dB gain, Binning off). The z-slices demonstrated consistent staining intensity through the depth of the spheroids (Supplementary Fig. 3). On Day 0, the spheroids, pre-stained with Hoechst during spheroid formation, were imaged live immediately after gelation of the collagen, and the spheroid locations were saved for use again during Day 2 imaging. On Day 2, the samples were fixed with 4% paraformaldehyde, permeabilized using 0.25% Triton-X 100 and stained again with Hoechst, at a 1:1000 dilution to obtain a strong fluorescent signal in the nuclei. They were further stained for F-actin visualization using Alexa Fluor® 488 phalloidin (Life Technologies) at a 1:100 dilution. The images of each z-stack were processed using the Keyence BZ-X800 Analyzer to create maximum z-projection images (Fig. 1A, Supplementary Fig. 4), increase the image contrast and convert them to grayscale.

**Figure 1:**
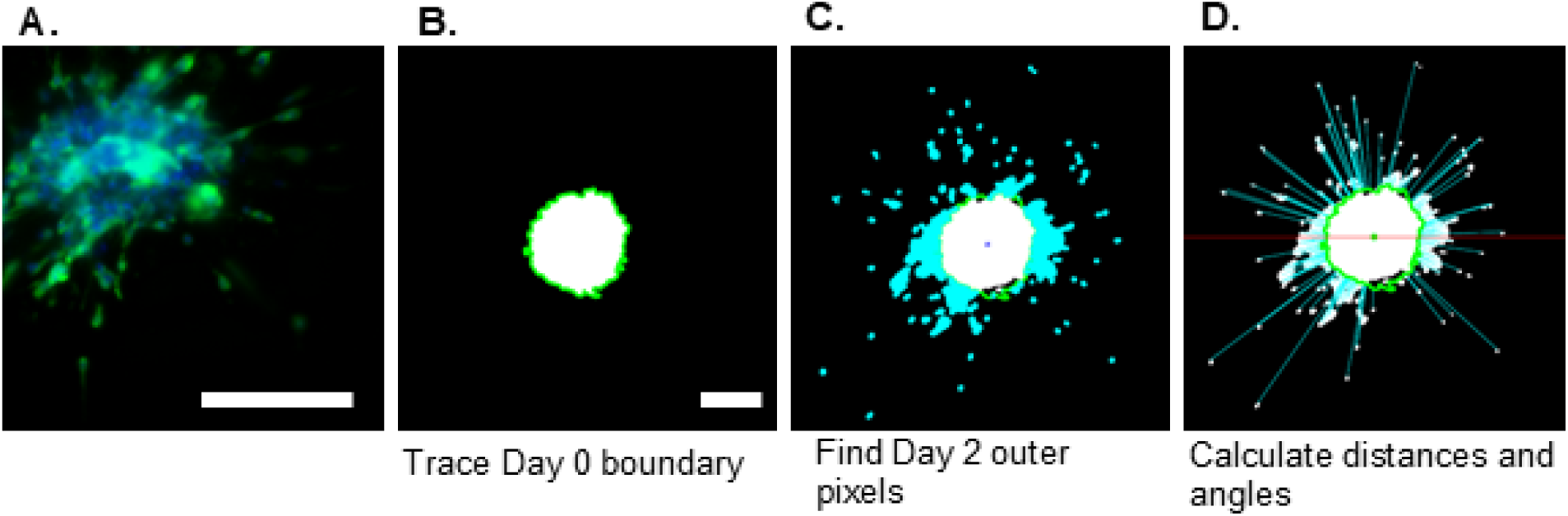
Z-stack image processing and quantification. Images were processed to create maximum z- projection images from each z-stack as demonstrated by a representative Day 2 image (A). The Day 0 boundary was segmented (B) and the pixels of the Day 2 image located past the boundary were identified (C). The distances and angles of the pixels outside of the boundary were calculated with reference to the spheroid boundary and center. For clarity, only a portion of the distance lines are shown (D). Scale bar: 200µm. Panel A: Representative spheroid is imaged at 20X magnification and off-set to highlight invading cells. Panels B-D: 10X magnification.

Because the spheroids are mixed into the collagen hydrogel solution, it was common to observe spheroids that were close to each other or out of the vertical field-of-view. Spheroids that were out of the field of view or too close together (less than 400 µm apart or with migration paths of different spheroids that could cross) were not imaged (Supplementary Fig. 5).

### Quantification of cell invasion and migration into matrix

Image quantification was carried out using a custom MATLAB program (ver R2024a, MathWorks®, https://www.mathworks.com/) using the image processing package. To provide users with the option of a “single-package tool” we also include binarization and data organization components to the program, although the quantification component is the focus of this study as it is the novel component. The program is divided into three distinct components, allowing users to integrate them with their preferred tools. (1) A semi-automatic binarization and image correction tool that also applies a circular mask around the images edges; (2) a fully automatic invasion quantification tool that computes invasion metrics given spheroid images from an initial and final timepoint; and (3) a data consolidation tool that compiles the data computed for each spheroid into one file for easier organization.

For image binarization, the grayscale images were contrast enhanced, background subtracted if needed, and binarized with a global threshold strategy, which is determined by the user to best represent the grayscale image. A global threshold of 0.16 was found to work best for most of our images from the input range [0, 1]. Some spheroids required further image correction, so for those images, remaining artifacts and noise were removed manually using our custom image correction tool. A circular mask was then automatically applied to the images to prevent biases towards the long axis and corners of the frame by excluding those pixels. These steps generate a set of Day 0 and Day 2 binarized images for each batch of images.

For automatic invasion quantification, the Day 0 spheroid edge was located and automatically segmented to determine the boundary of cell invasion (Fig. 1B). The centroids of the Day 0 and Day 2 spheroids were identified and used to overlap the Day 0 boundary and the Day 2 binarized image. From there, the locations of all the pixels of the Day 2 image that lay outside of the boundary were found (i.e., “the outer pixels”) (Fig. 1C), and the radial distances of each of these pixels from both the boundary and the spheroid centroid were calculated as well as their angles from the centroid, measured clockwise from the x-axis (as in a polar coordinate system) (Fig. 1D).

The change in spheroid area *(ΔA)* i.e., the stained pixel area outside of the initial spheroid boundary, is calculated by summing the number of pixels in the image and subtracting the Day 2 area to the Day 0 area, equation (1).

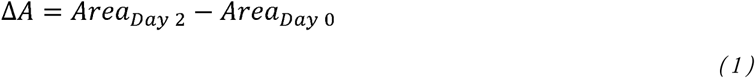

By locating the individual pixels in the images, rather than cell or nuclei centroids, pixel-level distance calculations can be performed which avoids the difficulty of distinguishing clustered cells. The radial distance *(d)* is calculated from the differences of the x and y distances from the boundary (x_b_, y_b_) and the outer pixels (x_p_, y_p_) along a radial line emanating from the centroid of the Day 0 image to the outer pixel, equation (2) (Fig. 1D):

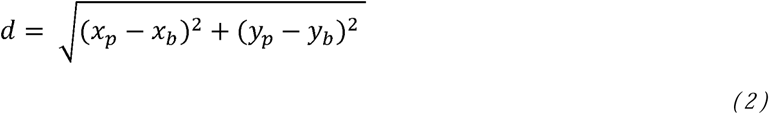

The angles of invasion are obtained by calculating the arctangent of the x and y distances to the spheroid centroid, (x_0_, y_0_), equation (3).

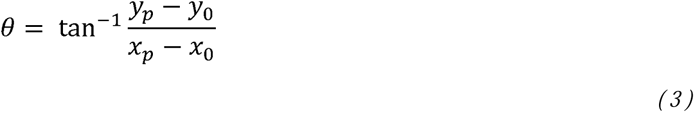

The invasion area moment of inertia *(I)* was evaluated as an integrative measure of the area change and the distance from the spheroid boundary. The area moment of inertia, also known as the second moment of area, is an established parameter in mechanics that reflects how the points of an area (dA) are distributed with regard to an arbitrary axis. The area moment of inertia is an important term in engineering; it describes the resistance of a body to bending or torsion, as mass further away from the neutral axis has a greater effect on the resistance. We feel that our analogous metric will be helpful for researchers by providing a parameter for overall invasiveness of a cell type or population that reflects how the points of nuclear area are distributed away from the initial spheroid boundary giving more weight to the cells that invade further into the matrix. The radial moment, *I_r_*, was calculated by multiplying the squared radial distances (*d*) of each outer pixel (*i*) to the boundary with the area of each pixel (i.e., the pixel size) *(dA)* and summing over all the outer pixels, equation (4).

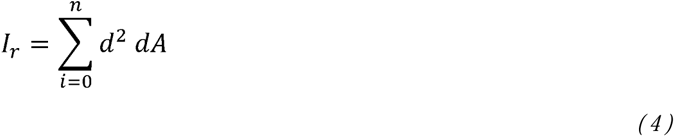

### Principal component analysis

In addition to being able to quantify the directional migration along the camera x- and y-axes, the invasion area moment of inertia can be calculated along any arbitrary axes (x’, y’) using coordinate transformation. Most notably, the directions of maximum and minimum invasion and extent of invasion in each direction can be expediently calculated using principal component analysis (PCA). Coordinate transformations were performed in MATLAB to reorient the distance values along the principal components (directions of maximum and minimum invasion). The extent of invasion is then determined by calculating the directional moments of inertia (*I_x_’* and *I_y_’*) along the principal axes, by summing the transformed directional distances (*x’* and *y’*), equation (5), and calculating the mean distance in the principal directions.

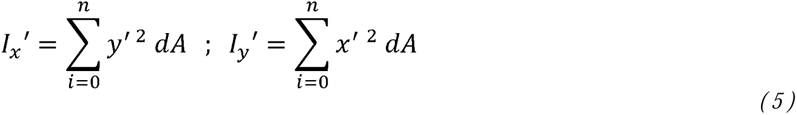

### Code availability

Our custom MATLAB image analysis program is shared on GitHub to facilitate use by other researchers. <https://github.com/rmungai/SpheroidInvasionAnalysis>

To expand user accessibility, we have published a Python version of the program as well as a downloadable GUI for users without computer programming backgrounds. <https://github.com/rogerh2/SpheroidInvasionAnalysis>

Detailed descriptions of the scripts are also included on the GitHub repositories.

### Statistical analysis

An outlier analysis was performed on the dataset using MATLAB by detecting and removing values more than three scaled median absolute deviations (MAD) from the median. The normality of the data set was assessed using MATLAB via the D’Agostino & Pearson omnibus normality test employing a significance level α= 0.05. After determination of normality, significant differences among groups were analyzed using the sjstats library in R using one-way analysis of variance (ANOVA) followed by Tukey’s Honest Significant Difference (HSD) post hoc test. Violin plots of the area change, mean distance from the boundary, and area moment of inertia per spheroid were generated using the ggplot library in R and were utilized to show the entire data distribution of all spheroids. Numbers of spheroids and biological replicates are provided in the figure captions.

## RESULTS

We formed multicellular spheroids of different cell types using non-adherent agarose molds (Supplementary Fig. 1), embedded them into a collagen hydrogel and observed the extent of cell invasion into the hydrogel after two days (Supplementary Fig. 2). The cells were stained for nuclei on both Day 0 and Day 2 to allow for comparison of invasion over the culture period (Supplementary Fig. 4). We then binarized and quantified the images using our custom MATLAB program which identifies the locations of all the pixels of nuclei past the Day spheroid boundary (Fig. 1) and calculates the area change over the culture duration and the distances of the outer pixels to the boundary as well as their angles. To determine if a segmentation-free analysis allows for superior image quantification of multicellular 3D spheroids than object-based analysis, we compared our method to a standard object-based approach using Cell Profiler (Supplementary Fig. 6). Nuclei that have dissociated from the spheroid mass are accurately segmented; however, the individual nuclei within the spheroid mass are not segmented as effectively as the dissociated nuclei (Supplementary Fig. 6D, E). In contrast, a segmentation-free approach allows for identification of all outer pixels irrespective of object classification (Supplementary Figure 6F). These results demonstrate the difficulty of distinguishing and segmenting individual cell nuclei when they are tightly clustered into multicellular spheroids and confirm that a segmentation-free approach is more advantageous in such cases.

Next, individual example spheroids were selected to demonstrate the nuances of the invasion metrics. We generated polar plots depicting invasion past the boundary for the selected spheroids and calculated their quantitative metrics (Fig. 2). As demonstrated by comparing polar plots of individual NHF and SMC spheroids (Fig. 2A&B), the area change is not necessarily correlated with distance. Panel B shows a spheroid that does not invade as far as the spheroid in Panel A, resulting in a lower distance, but the invasion distribution is denser, resulting in a higher area change. Due to these differences in invasion density and distance, these spheroids have a nearly equal value (∼0.001 mm^4^) for the radial area moment of inertia demonstrating that the moment metric is less influenced by invasion distribution differences than the area change and distance metrics alone.

**Figure 2:**
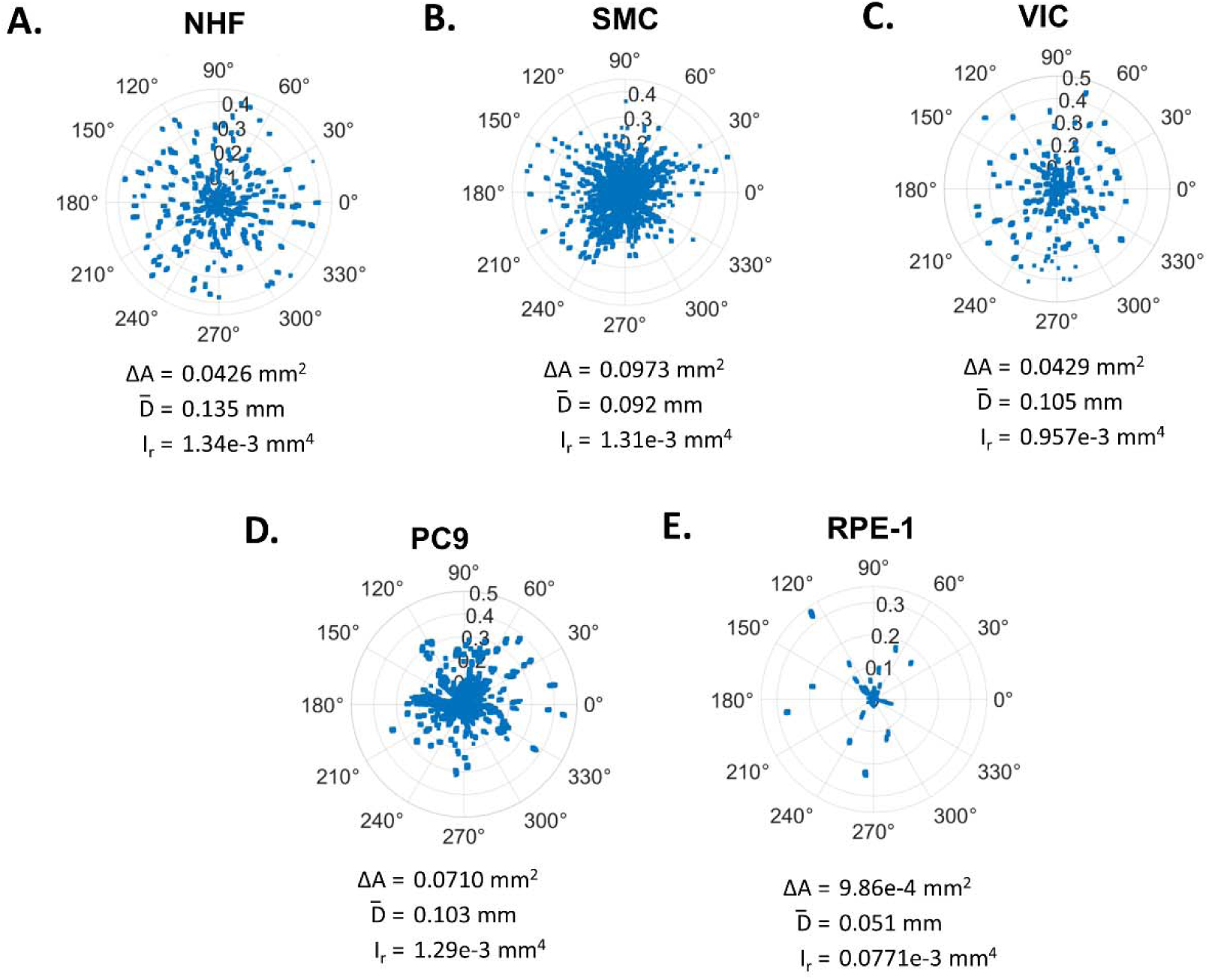
Visualization of variation of invasion behavior. Polar plots demonstrate invasion behavior of individual spheroids on Day 2. The distance (µm) versus angle is plotted for each pixel past the Day 0 spheroid boundary for an example VIC (A), NHF (B), SMC (C), RPE-1 (D) and PC9 (E) spheroid. The boundary is represented as the plot center (0,0). Invasion metrics for example spheroids, area change, ΔA, mean distance, 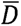, radial moment, I_r_, as defined in the text, are provided below each plot.

We also aimed to explore whether the degree of thresholding has a greater effect on the quantification metrics obtained via a segmentation-free or object-based quantification approach (Supplementary Fig. 7). Using MATLAB, both approaches were compared for the dissociated individual nuclei of a representative spheroid at low, medium and high global threshold values. As expected for both approaches, the quantification metrics of area, distance and moment increase as the threshold decreases. Interestingly, the number of single nuclei segmented in the object-based approach does not increase as the threshold decreases, rather, the relationship is non-monotonic indicating that segmentation-free analysis has a more predictable relationship with threshold than an object-based approach.

In addition, we compared the computation time for the automatic invasion quantification program for a batch of images. The object-based method is only slightly faster than our pixel-based method at 1.7 and 2.1minutes per image set (Day 0 and Day 2), respectively (see supplementary section for computer specifications).

### Directional invasion assessment

Next, we assessed whether the area moment of inertia can be used to assess invasion directionality resulting from a guidance cue, particularly in cases where directional invasion does not exclusively occur along the x- and y- axes. We performed principal coordinate analysis (PCA) to a representative spheroid exhibiting directional invasion due to its proximity to a stiff silicone post within the gel (Fig. 3A, right). The post introduces constraint in the hydrogel which causes directional invasion towards it via contact guidance [46,57–59]. A binarized image of the spheroid demonstrating directional invasion was corrected to mask the post from the image (Fig. 3C, right) and quantified as described above. PCA resulted in the determination of the principal angles of maximum and minimum cell invasion (Fig. 3D, right). A coordinate transformation was then performed to calculate the extent of invasion along the principal angles via the mean distance (0.104 mm and 0.071 mm) and the directional area moments of inertia (0.227e-3 mm^4^ and 0.103e-3 mm^4^). The fold change of mean distances and directional moments were calculated (by taking the ratio of the maximum to minimum directions) and found to be 1.5X and 2.2X greater in the direction of the post, respectively, thus indicating directional invasion. To validate the directional analysis method, a control spheroid (which was not located next to a post) was also quantified for comparison (Fig 3, left). The mean distances along the principal angles for the control spheroid were found to be 0.0984 and 0.0942 mm, and the directional moments were found to be 0.369e-3 mm^4^ and 0.329e-3 mm^4^ (Fig 3D, left) resulting in fold change values of 1.1X for both metrics. In comparison, the mean distances and directional moments fold change values are 1.4X and 2X greater for the spheroid with the post (Fig 3, right). Taken together, these results demonstrate that this method effectively detects and quantifies invasion directionality.

**Figure 3:**
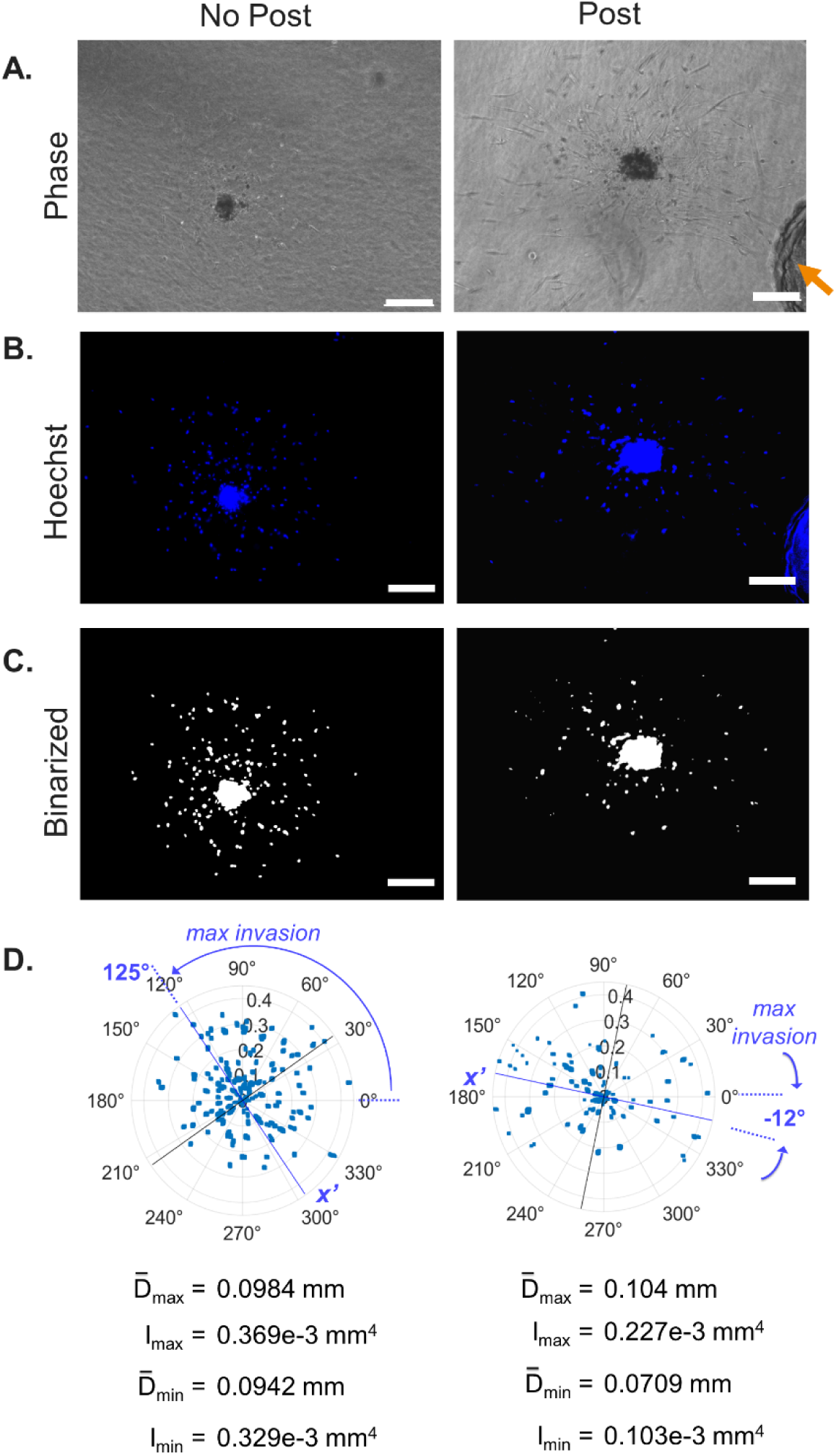
Quantification program also assesses invasion directionality. Directional invasion was demonstrated by imaging a collagen gel-embedded VIC spheroid growing near a structural post (indicated by arrow) of the culture well for four days. Phase, Hoechst-stained and binarized spheroid images on Day 2 (no post control) and Day 4 (spheroid with post). Scale bar: 200 µm (A-C). PCA of quantified data plotted as a polar plot centered at the mean. Principal angles are indicated (blue line: maximum invasion at 125° and −12°; black line: minimum invasion). Polar plot depicts the distance from the boundary (mm) versus angle for each pixel. Mean distances and directional moments along the principal angles were calculated to quantify invasion directionality and are provided below the polar plot (D).

### Comparison of spheroid invasion between cell types

We then aimed to compare invasion extents of the five cell types and found that, while all five cell types invaded into the collagen hydrogel, the extent of invasion varied substantially between the cell types (Fig. 4A, Supplementary Fig. 4). Qualitatively, the RPE-1 cells demonstrated the least invasion while the SMCs demonstrated the most invasion. The PC9 cells demonstrated the second-most invasion with the NHFs and VICs following, respectively.

**Figure 4:**
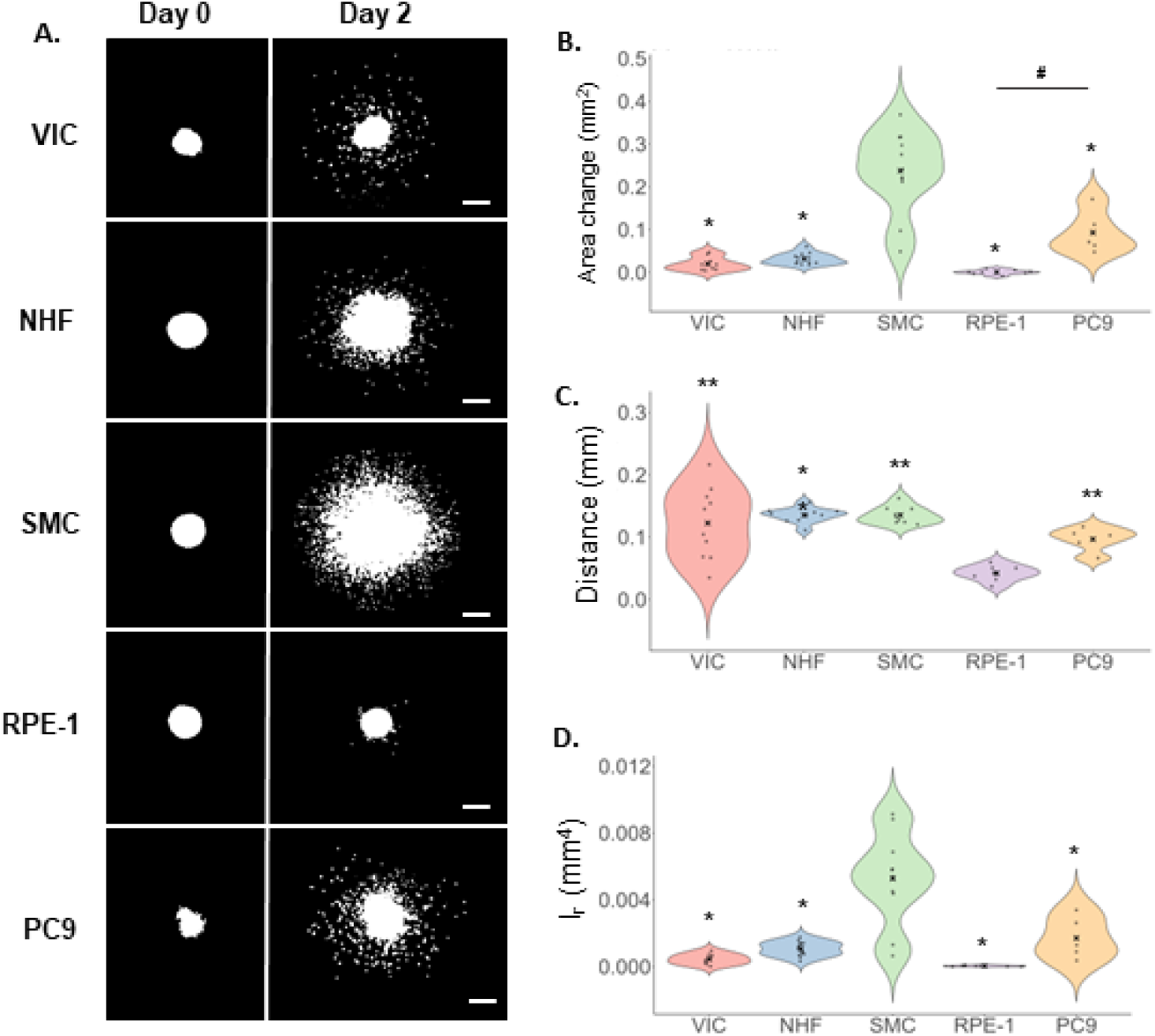
Invasion of cells from spheroids into collagen hydrogels. Representative images of invasion on Day 0 compared to Day 2 (A). Top to bottom: VIC, NHF, SMC, RPE-1 and PC9. Binarized maximum projection images of cell nuclei stained with Hoechst fluorescent dye at 10X magnification. Scale bar: 200 μm. Image quantification results for the area change (B), mean distance (C) and radial area moment of inertia (*I_r_*) (D) for VIC, NHF, SMC, PC9 and RPE-1 cells. Biological replicates are VIC=3, NHF & SMC=2, and RPE-1 & PC9=1. Number of spheroids are indicated by dots on each plot and are: VIC n=9, 10, 8 (A-C respectively), NHF n=12, 10, 13 (A-C respectively), SMC n=11, 9, 10 (A-C respectively), RPE n=8 (all), PC9 n=5 (all). “x” indicates the mean value. Significance at p<0.05 determined by one-way ANOVA with Tukey’s HSD post hoc test (exact p-values provided in Supplementary Table 1). * and ** indicate significant difference to SMC (B) and (D) and RPE-1 (C) respectively whil<u>e_ # _</u> indicates significant difference between the specified groups (B).

We then quantified the images to obtain the invasion metrics. The area change was found to be the highest for the SMCs and all other cell types had a significantly reduced area change in comparison indicating SMCs as the most invasive (Fig. 4B). The PC9 cells had the second-highest area change followed by the NHF, VIC, and RPE-1 cells respectively. The area change of the PC9 cells was significantly higher than that of the RPE-1 demonstrating that, as expected, the cancerous epithelial cell type is much more invasive than the wild-type epithelial cell type.

For the mean invasion distance, SMCs and NHFs were found to have the longest distance followed by VIC, PC9 and RPE-1 cells (Fig. 4C). All cells had a significantly higher mean distance than the RPE-1 cells, demonstrating that the epithelial cell type was, as expected, the least motile compared to the more migratory mesenchymal-type and cancer cells. SMCs did not demonstrate a significantly longer distance than other cells apart from RPE-1. This difference in the SMC results for area change and distance confirmed our previous finding that a high invasion area change does not necessarily indicate a longer invasion distance.

For the radial area moment of inertia of invasion (*I_r_*), the SMCs were found to have the highest value which was significantly higher than all the other cell types, as with area change (Fig. 4D). This finding suggests that cell proliferation may play a larger role in the SMCs, indicated by a higher number of pixels (i.e., higher area change). Since the area moment of inertia metric incorporates both invasion area and distance into the calculation, the moment calculation demonstrates that SMCs are the most invasive cell type overall, a finding that may have been overlooked if invasion distance was used alone.

## DISCUSSION

In this work, we develop an *in vitro* imaging and analysis method to objectively assess the extent of cell invasion from multicellular spheroids into a surrounding 3D hydrogel. Individual spheroids are imaged at initial and final time points allowing for accurate determination of cell invasion over the culture period and overcoming limitations of methods that only analyze the final state of invasion [34,39,42]. The projected distribution of nuclear pixels past the Day 0 boundary is automatically identified to calculate common invasion metrics, e.g., the increase in area and mean distance from the boundary, as well as the area moment of inertia, our new integrative metric which quantifies the overall extent of invasion. These metrics are calculated with reference to the initial boundary of the spheroid at Day 0 which is automatically segmented. We introduce coordinate transformation to identify the directions and extent of most and least invasion for cases where the invasion may not be radially symmetric due to directional mechanical or chemical signaling gradients. We demonstrate the utility of our method using five different cell types of varying invasive potentials and show that the metrics quantitatively capture the differences in invasiveness among the cell types.

By automatically segmenting the initial spheroid to generate the boundary of invasion, we lessen the effect that spheroid size differences can have on the invasion metrics. While this approach is not an all-encompassing solution to the challenge of generating consistently sized spheroids, as demonstrated by the variability in our invasion metric results (Figure 5), it aids in reducing the effect of spheroid size on invasion metrics as previously described [47]. This approach is especially useful for cell types that do not reliably form uniform spheroids (Supplementary Fig. 8) [30,38,60,61]. A limitation of this method is that it assumes that invasion occurs from cells located at the spheroid boundary, thus it does not account for the additional distance traveled by cells invading from deeper within the spheroid. However, this method provides a key advantage over the majority of invasion quantification methods that assume all spheroids are the same size [30,39,42,44,49]. Automated boundary segmentation makes spheroid invasion analysis more flexible for non-uniform spheroid sizes and shapes and, by extension, makes it easier to quantify spheroids of various cell types.

To our knowledge, we are the first to introduce a pixel-based method for invasion analysis. Using pixels avoids the difficulty of distinguishing individual cells from a cluster. Comparing our segmentation-free approach to a standard cell segmentation method (Cell Profiler ver 4.2.6, www.cellprofiler.org) [62], we demonstrate that cell segmentation is challenging for the 3D multicellular spheroid bulk which is why most spheroid quantification studies segment the spheroid bulk as one object [47,48,63–65] (Supplementary Fig. 6). In addition, manual tracing of cell boundaries is user-dependent and time consuming and cannot be completed if the cell boundaries are not evident [8,30,38–44,64]. Automated spheroid invasion quantification methods such as TASI segment the spheroid mass as one object and, as a result, are limited to assessing the distance of invasion from the spheroid centroid or core boundary to the edge of a dense invasion front [47,48]. This limitation is also present in machine learning approaches [66] as well as in methods that quantify morphological features for spheroids not embedded in ECM, such as AnaSP and INSIDIA [64,65]. Our method has the key advantage of accounting for all pixels past the boundary—a distinction especially important for cell types (such as primary cells) which separate into individual clumps and do not form a dense collective invasion front, as we have demonstrated in Figure 3 and 5. This advantage is especially notable when compared to our finding that pixel-based quantification is only 29 seconds slower per spheroid than a similar object-based quantification.

To determine the sensitivity of our pixel-based approach to thresholding, we analyzed the dissociated single nuclei invading from a representative spheroid at low, medium and high global threshold values and compared the results to a standard object-based method (Supplementary Fig. 7). As the threshold is decreased, an increasing number of pixels surpass the threshold value leading to larger area, distance and moment values for both the pixel-based and object-based methods at comparable degrees. We expected that the number of single nuclei identified in the object-based method would be less sensitive to threshold than the pixel-based metrics because changing the threshold value would simply change the sizes of binarized nuclei. However, we were surprised to find a non-monotonic relationship between the number of single nuclei and the threshold. The number of nuclei increased when the threshold was decreased from the high to medium values as expected, but the number of nuclei decreased from the medium and low threshold values due to nuclei either coalescing into the spheroid bulk or merging together into a single object too large to meet the size criteria for a single nucleus. Therefore, our pixel-based approach is not only comparable to an object-based approach for calculating invasion metrics but also potentially advantageous as it does not strictly report the “number” of cells or nuclei which, as we have demonstrated, has a less predictable relationship with threshold value.

Maximum projection z-stack images of fluorescently stained nuclei were utilized since the nuclei are high contrast and easy to binarize and can be easily stained for live imaging using inexpensive and readily available stains, (e.g., Hoechst) and are common for 3D spheroid invasion analysis [30,37,42,48,67]. If analysis of 3D invasion along the z-direction is needed, these methods can potentially be applied to individual z-stack images [68]. Further, quantifying cell nuclei rather than the cell body results in more accurate invasion area measurements because the cell nucleus, unlike the cell body, commonly resists large changes in shape during cell migration and invasion [53] due to being 5-10 times stiffer than the surrounding cytoskeleton [54]. This effect of cell shape on perceived cell size in 2D images is an issue overlooked by studies that assume invading cells are all the same size [8,30,34,37,38,44,49]. Thus, utilizing individual pixels of fluorescently stained nuclei is a powerful approach to quantifying spheroid cell invasion.

We introduce the application of the classical area moment of inertia parameter, which reflects how the points of an area are distributed with regard to an arbitrary axis and apply it towards spheroid invasion as an integrative “invasiveness” metric encompassing both the projected distance and area of invasion. Thus, the area moment of inertia quantifies the full extent of invasion. While the area change metric is useful for representing the change in cell number over the culture period [30,37–39,44], it may be skewed by variations in cell proliferation rates among cell types. The mean distance does not have this limitation, but it does not provide insight into the cell number [42]. Liu *et al.* use a similar metric combining cell area and distance of invasion (not squared as for the area moment of inertia) [30]; however, their method requires manual image tracing of cells from a uniform spherical bead whereas our metric is implemented more automatically and can be applied to common, arbitrarily shaped spheroids. Using specific cases, we demonstrate that differences in cell invasion distributions can skew the perception of invasion when using the area or distance metrics. For example, a spheroid with a dense invasion front with a high area change but a short invasion distance (Figure 3B) could be perceived as more or less invasive to a spheroid with a sparser invasion front with a lower area change but a longer invasion distance (Figure 3A). The difference in area values could be largely due to a higher level of cell proliferation for the SMC spheroid; the predominant mechanisms of invasion for each cell type can be explored in future work. Despite the differences in area and distance values, these spheroids undergo invasion to the same extent, as determined by the area moment of inertia (∼1.3e-3 mm^4^) which we propose is the best metric for quantifying invasion (c.f., panels A and B in Fig. 3). Therefore, the standard metrics of area change and mean distance, along with polar plots are best used to complement to the moment of inertia to explore the nuances between the invasion distribution differences.

Quantifying invasion directionality for cell populations is important for determining the effects of spatial gradients of chemical and mechanical factors. Most previous studies either track single cell directional migration [8,39,40,46,69] or quantify invasion of a population of cells along one direction [46,47]. We directly calculate the extent of invasion along x- and y-image axes using the area moment of inertia (*I_X_* and I*_y_*) as well as estimate it using the mean distances in the x- and y-directions. For simpler comparison, the extent of anisotropic invasion between cell types can be calculated by the fold change of the directional moments and distances. Ibrahim *et al.* similarly calculate the mean distance in the maximum invasion direction (by fitting it to an ellipse) [47]; however, their approach limits analysis to only the contiguous portion of the spheroid and omits the isolated cells whereas our approach includes all invaded cells. For greater generalizability, we also introduce the use of coordinate transformation to account for cases when invasion does not occur strictly along the axes of the image. We showed that calculating the mean distances and directional moments along the axes of maximum and minimum invasion using PCA is a straight-forward way to characterize the extent of directionality of invasion. As a demonstration, we applied the method to a spheroid cultured near a stiff silicone post which increases cell invasion along its direction via contact guidance and found the invasion (as measured by area moment of inertia) to be 2.2-fold higher in the direction of the post. This method can be used to assess the directional invasion response to guidance cues applied in any direction (e.g., mechanical constraints and chemical gradients); to our knowledge, we are the first to apply PCA to cell invasion analysis.

To expand user accessibility to our computer program, we have published an open-source graphical user interface for researchers without training in computer programming. In addition, the source code is also published in both Python (which is open-source) and MATLAB to allow for custom adaptation as desired and to cater to the researcher’s programming language preference. Taken together, these options broaden the ease of use of this spheroid invasion quantification method to biomedical researchers with a broad array of backgrounds.

## CONCLUSIONS

Here we develop a high-throughput objective quantification method of cell invasion from multicellular spheroids into a 3D extracellular matrix and demonstrate its utility with different metrics of invasion applied to cell types of various invasive potentials. The program is divided into three components to allow users to integrate them with their preferred tools; the main novel component is our fully automatic invasion quantification tool. Our pixel-based method can be applied to a variety of spheroid invasion studies, such as exploring the roles of ECM components and guidance cues (e.g., chemokines and dynamic stretch) in cell invasion. The use of a nuclear stain minimizes the effect of cell size variations, while live-staining the Day 0 spheroid allows for objective spheroid segmentation. Calculating the distances of all pixels past the Day 0 boundary, which is unique to each spheroid, minimizes the effects of spheroid size and shape on the calculated metrics of invasion. Thus, this automated custom image analysis tool is complementary to traditional segmentation methods and is a considerable addition to the field. We also introduce the area moment of inertia as an integrative metric of cell invasion. The program also utilizes coordinate transformation to allow researchers to assess invasion directionality in response to a guidance cue. This innovative approach to 3D cell invasion analysis has the potential to advance research in fields including wound healing, cancer metastasis, and the repopulation of decellularized tissue engineered scaffolds.

## DATA AVAILABILITY

The datasets used and/or analyzed during the current study are available from the corresponding author on reasonable request and are also available at the Harvard Dataverse using the following link: <https://doi.org/10.7910/DVN/VQH0BK>

## ACKNOWLEDGEMENTS

The assistance provided by Jamie Baines in conducting experiments is gratefully acknowledged.

This work was supported by the National Science Foundation (NSF) [CMMI 1761432], the National Institutes of Health (NIH) [1R15HL167235-01] and the American Heart Association (AHA) [20AIREA35120448]. The funders had no role in study design, data collection and analysis, decision to publish, or manuscript preparation.

## AUTHOR INFORMATION

### Contributions

K.L.B. and R.W.M. conceived the experiment which was performed by R.W.M., G.E.J., and K.W.P.. Results analysis was performed by R.W.M. who also quantified the data using MATLAB. R.J.H.II translated the quantification methods to Python and designed the GUI. R.W.M. wrote the manuscript draft which was edited by K.L.B. All authors approved the final version of the manuscript.

## ETHICS DECLARATIONS

### Competing interests

The authors declare no competing financial interests.

## Notes

https://doi.org/10.7910/DVN/VQH0BK

https://github.com/rogerh2/SpheroidInvasionAnalysis

https://github.com/rmungai/SpheroidInvasionAnalysis

### Competing Interest Statement

The authors have declared no competing interest.

### Summary of Updates

Clarified definition of "invasion" and the "area moment of inertia"; Figures 1 and 5 combined into single figure; wording of manuscript made more fluid

