## Supplemental Document for "Towards a More Objective and High-throughput Spheroid Invasion Assay Quantification Method"

*Correspondence to:

Kristen L. Billiar

Biomedical Engineering Department

Worcester Polytechnic Institute

100 Institute Road

Worcester, MA 01609

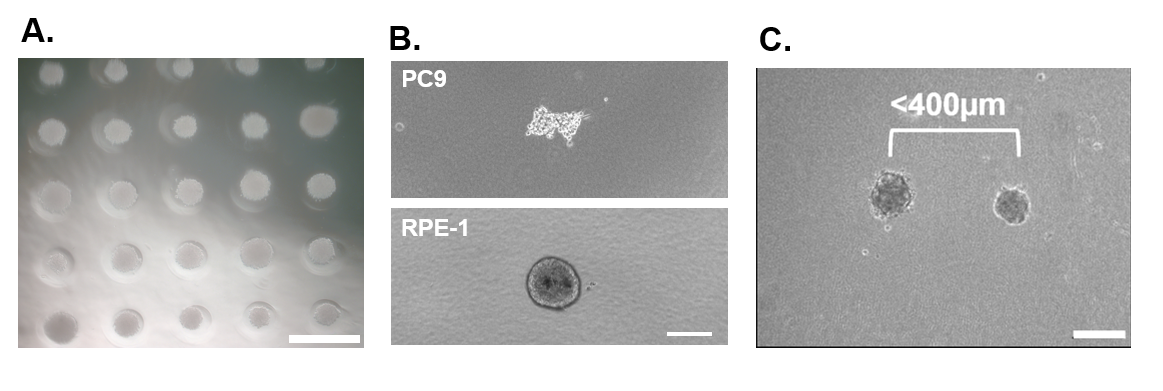

**Supplementary Figure 1:** Agarose molds form cell spheroids. Representative image of SMC spheroids. Scalebar = 1mm.

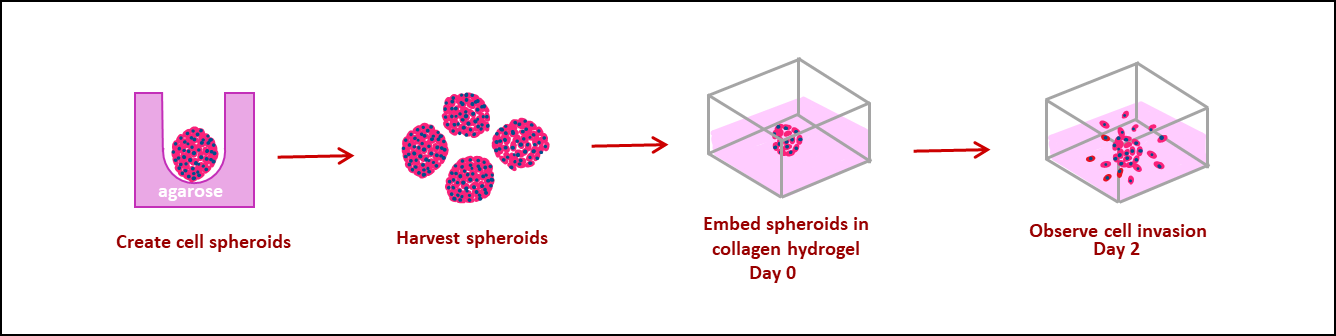

**Supplementary Figure 2:** Flow diagram of spheroid generation and gel embedding method. Cell suspensions within non-adherent agarose molds compact into dense spheroids. The spheroids are harvested the next day and embedded into a 2.0mg/mL collagen hydrogel and allowed to culture for two days after which the extent of invasion is observed.

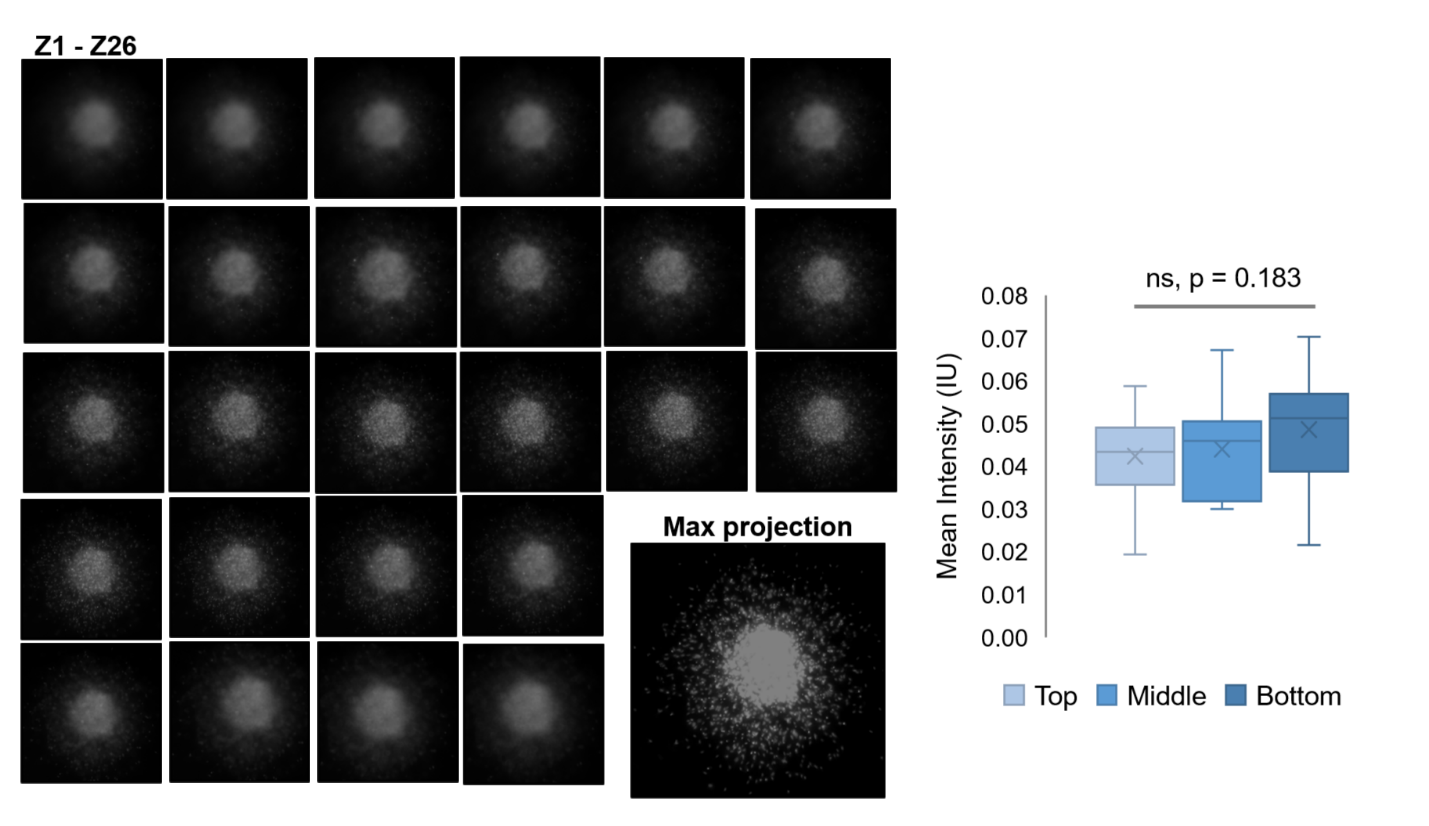

**Supplementary Figure 3:** Hoechst dye stain intensity is constant through depth of spheroids. Individual z-slice images (from top to bottom of spheroid, denoted Z1-Z26) and the corresponding maximum projection image for a representative SMC spheroid (left). Each image slice has a thickness of 10 µm. Image quantification of nuclei in top, middle and bottom z-slices (right). ns, indicates no significant difference among means as determined by one-way ANOVA (significance at p<0.05). n = 20 per region obtained for 10 spheroids with 2 nuclei per region. Plot generation and ANOVA performed in Excel. The normality of the data set was assessed using MATLAB via the D’Agostino & Pearson omnibus normality test employing a significance level α= 0.05.

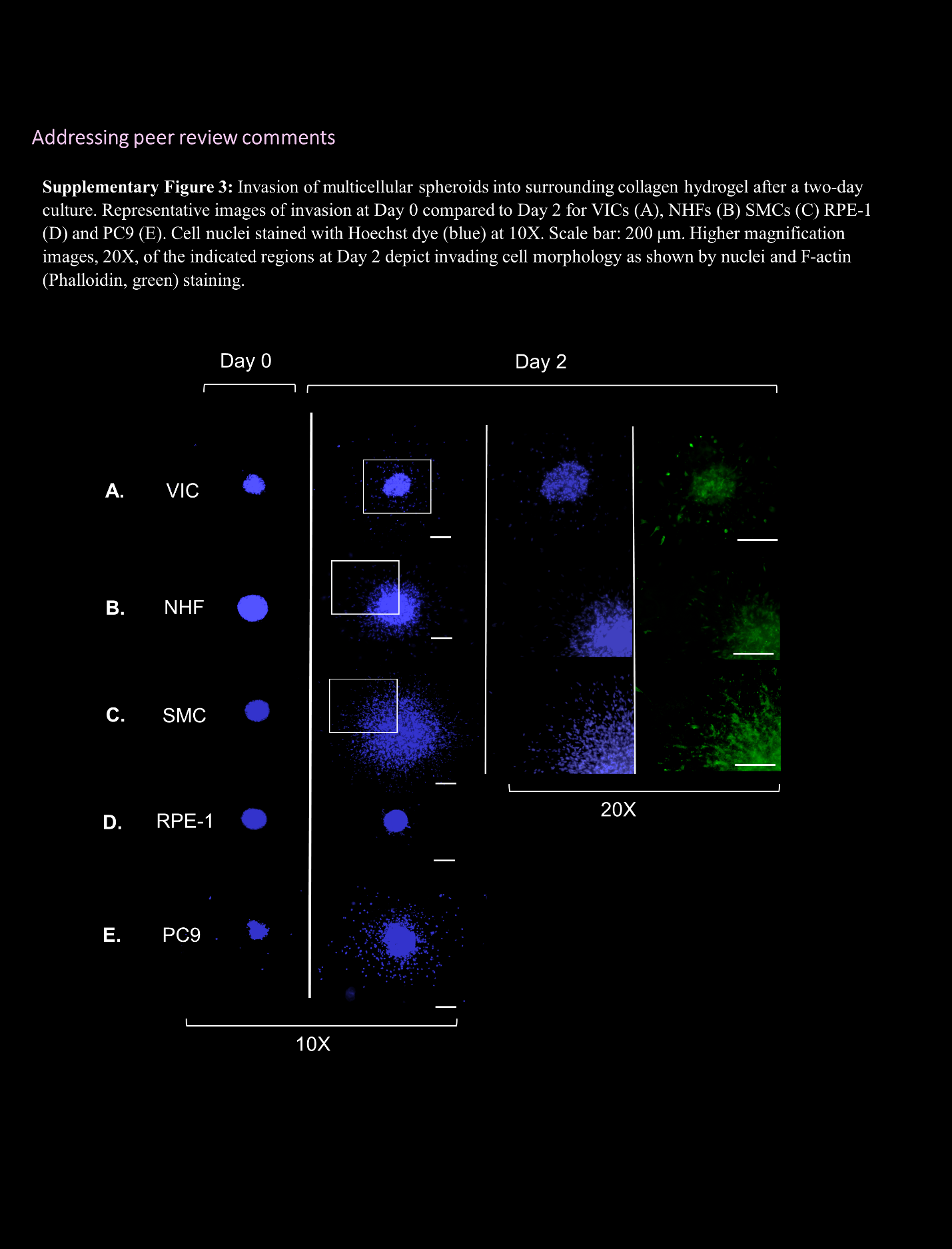

**Supplementary Figure 4:** Invasion of multicellular spheroids into surrounding collagen hydrogel after a two-day culture. Representative maximum projection images of invasion at Day 0 compared to Day 2 for VICs (A), NHFs (B) SMCs (C) RPE-1 (D) and PC9 (E). Cell nuclei stained with Hoechst dye (blue) at 10X. Higher magnification images, 20X, of the indicated regions (insets) at Day 2 depict invading cell morphology as shown by nuclei and F-actin (Phalloidin, green) staining (not done for RPE and PC9 cells). Scale: 200µm.

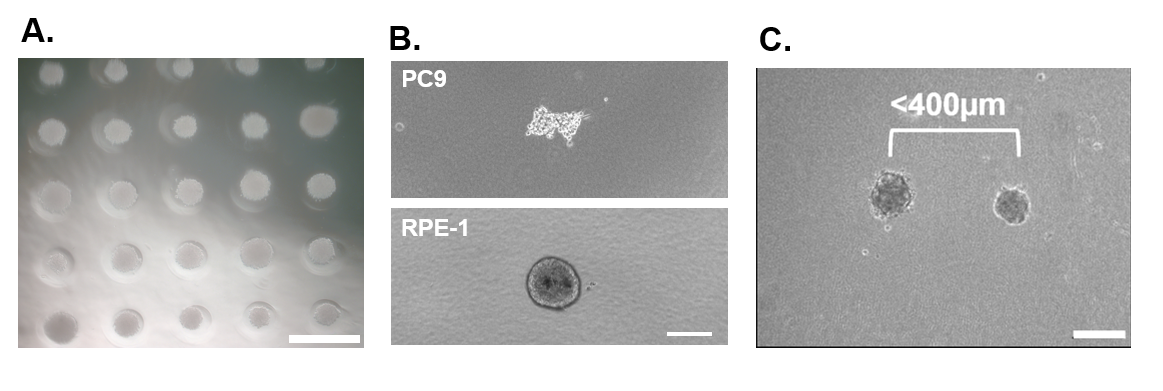

**Supplementary Figure 5:** Spheroids within 400µm of each other are omitted from imaging. Scale: 200µm.

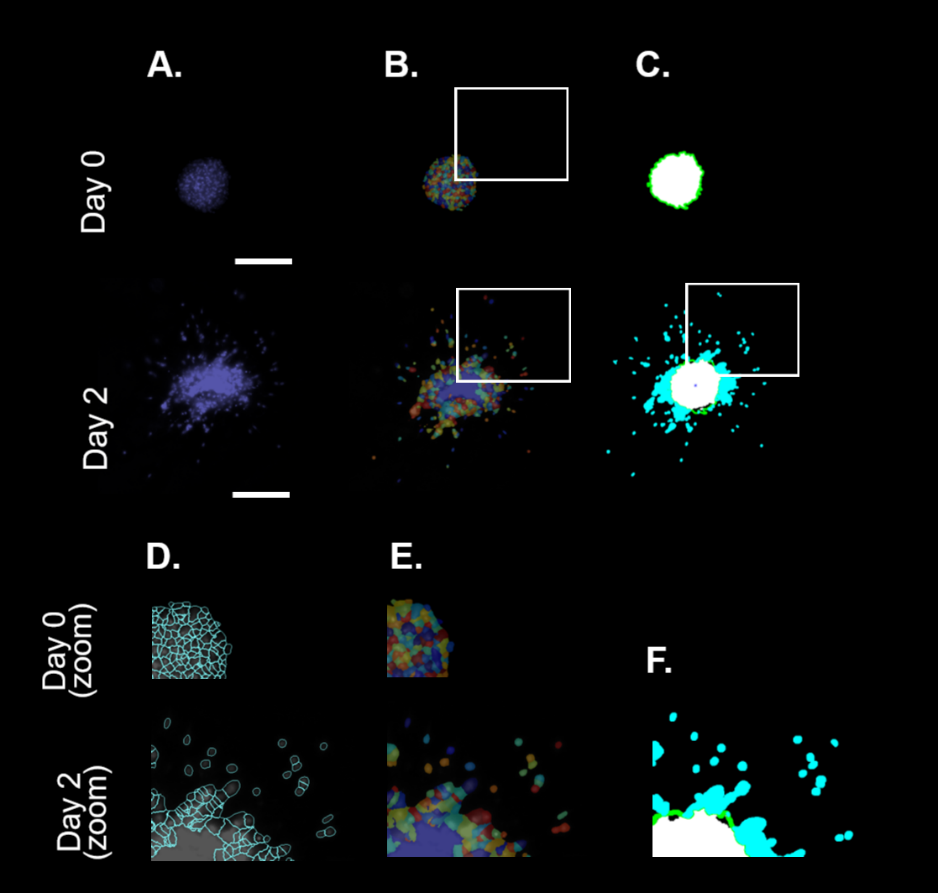

**Supplementary Figure 6:** Nuclei segmentation using Cell Profiler of multicellular spheroids invading a surrounding collagen hydrogel after a two-day culture. Representative images of Hoechst signal at Day 0 compared to Day 2 for VICs. Scale: 200 µm. (A) Depiction of cell segmentation using Cell Profiler (B) with inset depicting zoomed-in images of the segmented nuclei (E) and their outlines (D). The default module settings (global threshold strategy, minimum cross-entropy thresholding method) of the IdentifyPrimaryObjects module were utilized to segment nuclei. A size range of 3-20 pixels was used to determine the size of single nuclei. Comparison Day 0 and Day 2 image of MATLAB pixel-based analysis (C) with inset depicting the same zoomed-in region (F). Pixels outside of the Day 0 boundary (green) are depicted in blue.

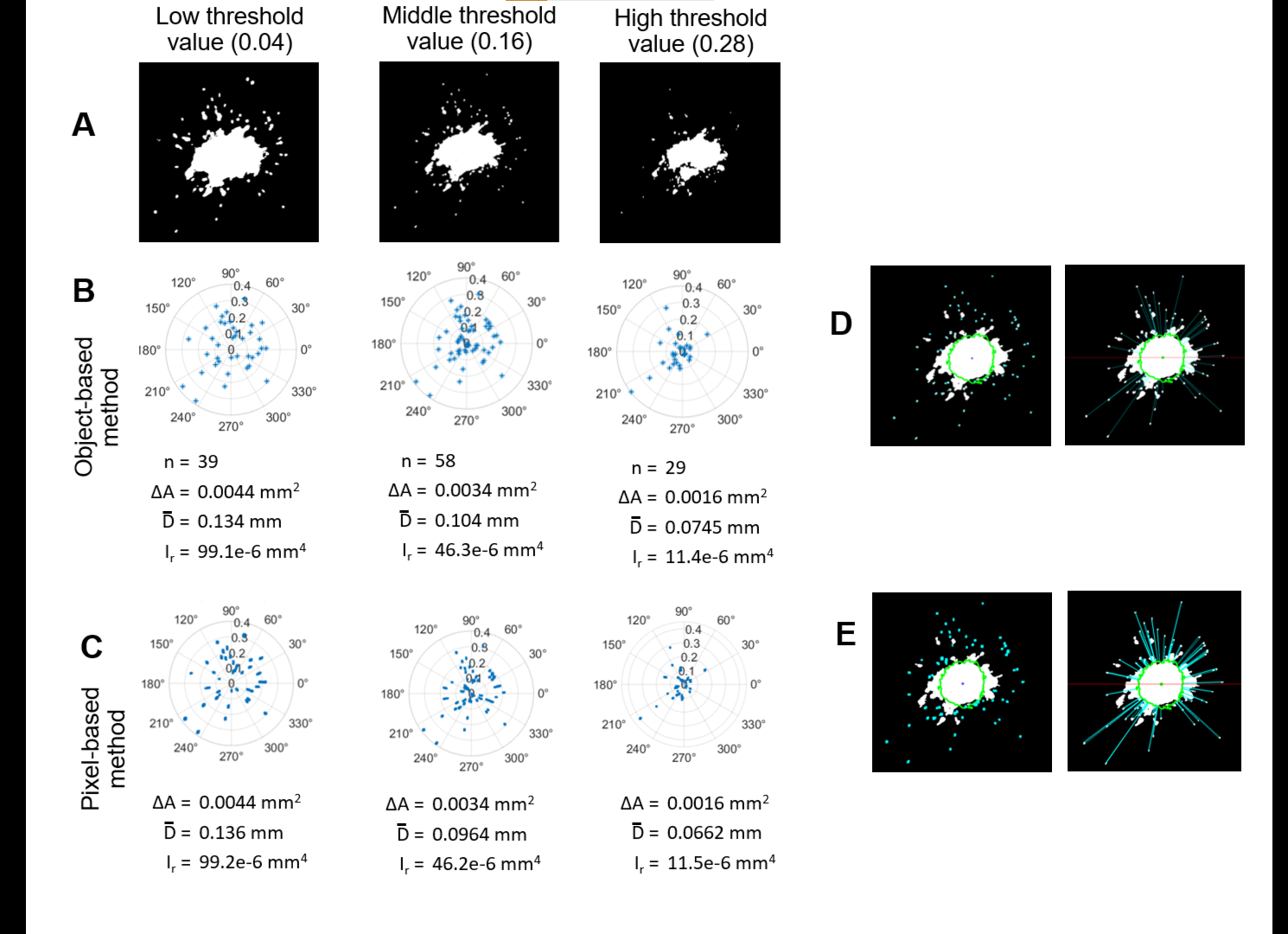

**Supplementary Figure 7:** Binarized images of a representative VIC spheroid using the different global threshold values in MATLAB: 0.04, 0.16, 0.28 (A). Polar plots depicting spheroid invasion at Day 2. The distance (µm) versus angle is plotted for each centroid (B) and pixel (C) past the Day 0 spheroid boundary for dissociated single nuclei. The boundary is represented as the plot center (0,0). Invasion metrics for the representative spheroid, number of single nuclei, *n*, area change, ΔA, mean distance, $\bar{D}$, radial moment, $I_{r}$, as defined in the text, are provided below the plots. Demonstration of centroid (D) and pixel (E) identification of the single outer nuclei and calculation of distances from the boundary (0.16 threshold value depicted). A size range of 30-300 pixels^2^ was used to determine the size of single nuclei.

The computation time of the MATLAB scripts were tested on a laptop computer running Windows 10 Home with the following specifications:

Processor	Intel(R) Core(TM) i5-8250U CPU @ 1.60GHz	1.80 GHz

Installed RAM	8.00 GB

System type	64-bit operating system, x64-based processor

Processing speed will vary with the user’s computer specifications.

**Supplementary Table 1:** Detailed table of p-values from the quantitative metrics of invasion results shown in Figure 5. P-values obtained by one-way ANOVA with Tukey’s HSD post hoc test.

| **Groups** | **Area change** | **Mean distance** | **Radial moment of inertia** |
| --- | --- | --- | --- |
| NHF - VIC | 0.984 | 0.888 | 0.869 |
| SMC - VIC | <0.001 | 0.896 | <0.001 |
| RPE - VIC | 0.935 | <0.001 | 0.977 |
| PC9 - VIC | 0.0986 | 0.546 | 0.533 |
| SMC - NHF | <0.001 | 1.000 | <0.001 |
| RPE - NHF | 0.668 | <0.001 | 0.498 |
| PC9 - NHF | 0.187 | 0.172 | 0.909 |
| RPE - SMC | <0.001 | <0.001 | <0.001 |
| PC9 - SMC | <0.001 | 0.185 | <0.001 |
| PC9 - RPE | 0.023 | 0.029 | 0.255 |

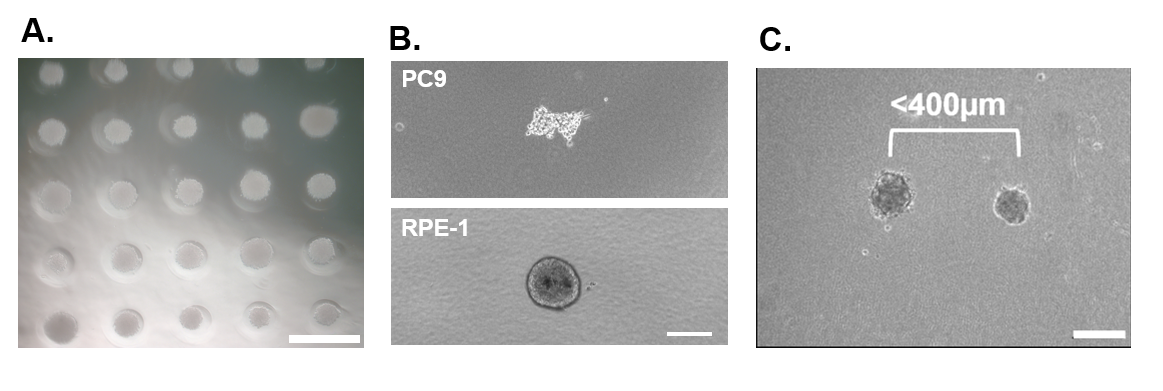

**Supplementary Figure 8:** Spheroid formation was not always uniform. Representative images of PC9 cell spheroids (top panel) which were the most non-uniform and RPE-1 cell spheroid which were more uniform. Scale: 200µm.
